# Using sentinel nodes to evaluate changing connectivity in a protected areas network

**DOI:** 10.1101/2023.04.25.538164

**Authors:** Paul O’Brien, Natasha Carr, Jeff Bowman

## Abstract

It has been recognized that well-connected networks of protected areas are needed to halt the continued loss of global biodiversity. The recently signed Kunming-Montreal biodiversity agreement commits countries to protecting 30% of terrestrial lands in well-connected networks of protected areas by 2030. To meet these ambitious targets, land-use planners and conservation practitioners will require tools to identify areas important for connectivity and track future changes. In this study we present methods using circuit theoretic models with a subset of sentinel park nodes to evaluate connectivity for a protected areas network. We assigned a lower cost to natural areas within protected areas, under the assumption that animal movement within parks should be less costly given the regulation of activities. We found that by using mean pairwise effective resistance (MPER) as an indicator of overall network connectivity, we were able to detect changes in a parks network in response to simulated land-use changes. As expected, MPER increased with the addition of high-cost developments and decreased with the addition of new, low-cost protected areas. We tested our sentinel node method by evaluating connectivity for the protected areas network in the province of Ontario, Canada. Our method can help provide protected areas ecologists and planners with baseline estimates of connectivity for a given protected areas network and an indicator that can be used to track changes in connectivity in the future.

## 1. Introduction

Protected and conserved areas are considered a fundamental conservation strategy for protecting biodiversity (McNeely, 1994; Naughton-Treves et al., 2005; Watson et al., 2014). With proper placement and management, protected areas can reduce species extinction rates, habitat destruction, and hunting mortality (Andam et al., 2008; Butchart et al., 2012; Hilborn et al., 2006), while also having the potential to secure valuable ecosystem services (Mitchell et al., 2021; Naidoo et al., 2008; O’Brien et al., 2023) and benefit human wellbeing (Naidoo et al., 2019). Nevertheless, recent reviews have called to question the effectiveness of current protected areas at reducing human pressure and biodiversity loss (Geldmann et al., 2019; Maxwell et al., 2020). Indeed, biodiversity inside and outside of park boundaries is increasingly threatened by accelerated human impacts including development, hunting pressure, and recreation (Barrueto et al., 2022; Craigie et al., 2010; Laurance et al., 2012; Tucker et al., 2018). Further, individual protected areas are often too small and isolated to support populations of large or vagile animals (Williams et al., 2022). Habitat loss, fragmentation, and degradation continue to be major threats to biodiversity loss through reductions in functional connectivity (Goodwin & Fahrig, 2002; Haddad et al., 2015) and can lead to isolation of populations within protected areas (Sawaya et al., 2019). While protected areas can help to safeguard biodiversity (Gray et al., 2016), human impact outside of parks can inhibit species ability to move (Tucker et al., 2018) and many species are expected to experience shifts in the location of suitable habitat with climate change, necessitating the need to move (Bellard et al., 2012; Chen et al., 2011; Parmesan, 2006). Therefore, maintaining and restoring connectivity among protected areas is a high priority for the protection of biodiversity within protected areas (Newmark et al., 2023).

Landscape connectivity is the degree to which landscapes facilitate or impede movement of organisms among resource patches (Taylor et al., 1993; Tischendorf & Fahrig, 2000). Connectivity is critical for species movement and gene flow through dispersal and migratory movements (Noss et al., 2012). Consequently, loss of connectivity can be a driver of species extinctions (Hooftman et al., 2016; Thompson et al., 2017) as isolated populations face increased risk of extinction due to inbreeding depression, stochastic events, and reduced opportunity for genetic rescue (Hoffmann et al., 2021; Pimm et al., 2006). Compared to isolated core areas, ecological networks, that is, networks of core areas (e.g., protected areas, OECMs, and unprotected Key Biodiversity Areas) and corridors are better able to connect populations, maintain ecosystem function, and support species persistence in the face of climate change and landscape modification (Heller & Zavaleta, 2009; Hilty et al., 2020; Schloss et al., 2022). Connectivity is therefore thought to be essential for the long-term persistence of biodiversity (Ward et al., 2020). While the need to build well-connected systems of protected areas has been identified as important by previous international biodiversity agreements (Aichi Targets) and more recently reinforced in the COP15 biodiversity agreement (Goal A and Target 3 of Kunming-Montreal Global Biodiversity Agreement; Convention on Biological Diversity, 2022), connectivity is often not incorporated into conservation planning (Carroll & Ray, 2021; Maxwell et al., 2020). To better address connectivity in conservation strategies, and more specifically into parks development and planning, we suggest that land-use planners and decision-makers across various jurisdictions (local to national) require the tools to (1) map contemporary connectivity among protected areas to guide identification of new areas for protection or restoration; and (2) evaluate the effects of future land-use changes or conservation interventions on connectivity of the protected areas network.

The rarity of connectivity being integrated into protected areas planning is not due to a lack of the methods, as a wide variety of techniques exist for assessing protected area connectivity. Circuit-theoretic models of connectivity are one such method that has gained increasing popularity. These models, which draw on the analogy between animals moving through a landscape and electrical current moving through a circuit, allow for the identification of multiple movement pathways between defined source and destination nodes (McRae et al., 2008). Typically, these nodes have represented core habitat patches that animals are likely to move among, often including protected areas. Many studies have used a circuit-theoretic approach with Circuitscape software to model connectivity among protected areas at various scales, including regionally (Belote et al., 2016; Dickson et al., 2017), on a continental scale (Barnett & Belote, 2021), and globally (Brennan et al., 2022). One output of Circuitscape is a cumulative current density map, measured in amperes, where current density is proportional to the probability of an animal using any given pixel during a random walk through the landscape. The resulting current density map can be used to identify areas that contribute importantly to connectivity between protected areas or other habitat patches or likewise, areas where connectivity is poor and could be restored. Circuit theoretic models provide a suitable framework for examining contemporary protected areas connectivity, which satisfies our first suggested requirement of tools for incorporating connectivity into conservation planning. The second requirement is being able to track future changes in connectivity through time, which may be achieved with use of a connectivity indicator.

Indicators are goal-specific metrics that can be measured to determine whether objectives are being achieved. In the context of connectivity, many different metrics have been developed (see Keeley et al., 2021 for a detailed review), however minimal progress has been made towards integrating these metrics into on-the-ground action (Theobald et al., 2022). Building on work by Jaeger (2000), Theobald et al. (2022) describe the desirable properties of a connectivity indicator including that it should reflect within- and between-patch connectivity, be computationally efficient, and be simple to measure and communicate. To this list, we add that a connectivity indicator should be repeatable to track progress in connectivity goals through time. Jaeger (2000) introduced a quantitative measure of landscape fragmentation, the effective mesh size (M_eff_), which can be interpreted as the probability of two animals, placed in different locations in a study area, encountering each other in the same patch. The effective mesh size satisfies many of the desirable properties of a connectivity metric and can be modified to incorporate within-patch connectivity (Deslauriers et al., 2018; Spanowicz & Jaeger, 2019); however, it may not be suitable for an analysis specific to protected areas and their surroundings, since M_eff_ would exclude unprotected natural areas within the matrix from measurements of connectivity. A more appropriate indicator of protected areas connectivity may be provided directly from Circuitscape.

Pairwise effective resistance is a second output of circuit theory models, and is a measure of the cost of moving between two nodes or habitat patches, calculated for all pairs of nodes. The more potential pathways there are between two nodes, the lower the effective resistance will be. Brennan et al. (2022) used a version of effective resistance, which they called the ‘Protected Area Isolation’ (PAI) index, to measure connectivity for a global protected areas network. The PAI index calculates the cost of movement between a focal protected area and all other protected areas in the network, and so reflects the degree of isolation of the focal park. This metric adds to existing global connectivity metrics by incorporating functional connectivity and can be implemented efficiently at a global scale, however, it may be less suitable when the objective is to track network connectivity through time given that calculation of the PAI index is dependent on the membership of the contemporaneous protected areas network. Further, because the number of pixels substantially increases at finer resolutions (< 1 km), which may be more useful to regional planners, calculation of the PAI index may not be as computationally efficient owing to the need to run Circuitscape twice – once to produce the current density map and again to calculate the PAI index using a different mode (Brennan et al., 2022). We suggest an alternative would be to use the mean pairwise effective resistance (MPER) as an indicator of connectivity. This is the mean of all pairwise effective resistances and reflects a measure of overall network connectivity. Further, both pairwise effective resistance and a cumulative current density map are produced within a single run of pairwise mode Circuitscape, making this a more efficient method.

One consideration with our proposed MPER indicator, and one that arises with other connectivity indicators, is that they are not necessarily repeatable across time if the set of network nodes is dynamic. That is, as the network changes through addition or loss of protected areas, the set of nodes changes and so recalculations of the indicator across time are no longer comparable (i.e., there is a conflation between space and time). To deal with this problem, we propose a modified version of park-to-park connectivity methods whereby we model connectivity using a set of sentinel nodes. We define sentinel nodes as a subset of protected areas used to assess connectivity of the protected area network over time. This fixed set of nodes provides a repeatable framework for modelling connectivity and tracking changes in connectivity of a network through time. To develop the idea of a sentinel node indicator, we first carried out sensitivity analyses to determine (1) how well a subset of nodes captures connectivity of a full park network; (2) the ability of the method to detect changes in the network; and (3) how many sentinel nodes are required to produce an accurate estimate of connectivity for the full network. We conducted these tests on smaller, but real-world landscapes using the 300-m resolution cost surface produced by Pither et al. (2023). Next, we tested the ability of our method at a larger scale by modelling connectivity of the protected areas network in the province of Ontario, Canada. The current protected and conserved areas network in Ontario consists of >1400 protected areas covering 10.9% of Ontario (Environment and Climate Change Canada, 2023) and accounting for 1.17% of the national coverage (Environment and Climate Change Canada, 2023). Canada has committed to protecting 30% of terrestrial lands in a well-connected network of protected areas by 2030. The current state of connectivity of the Ontario protected areas network is unknown. Our assessment will help to provide a baseline understanding of park-to-park connectivity for the province’s protected areas, and an indicator of parks connectivity for future use. Our analysis will also support decisions required to achieve ambitious national targets.

## 2. Methods

### 2.1. Cost Surface and Nodes

We modelled landscape connectivity using a circuit theoretic approach, which takes advantage of the analogy between electricity moving through a circuit and animals moving across a landscape, allowing for the identification of multiple movement pathways. Electricity travels through a circuit according to a random walk, and animals often move using a random walk, so circuits can be used to model these movements (Doyle & Snell, 1984). Modelling connectivity using circuit theory requires two inputs: a cost surface and a file with the location of source and destination nodes. A movement cost surface represents features in a landscape (e.g., roads, rivers, forests, etc.) by the degree to which they facilitate or impede movement (McRae et al., 2008). For the following analyses, we used the 300-m resolution cost surface of Canada produced by Pither et al. (2023), which classifies the landscape according to a cost scheme using 4 values including 1 (low cost), 10, 100, and 1000 (high cost) (Table 1). Bowman et al. (2020) showed that current density maps are not sensitive to the absolute costs of a cost surface provided that costs are in the correct rank order. Pither et al. (2023) modelled landscape connectivity for terrestrial non-volant fauna, such that natural features like forests, wetlands, and grasslands were assigned a low cost, while anthropogenic features like roads and cities, as well as large water bodies and mountains were represented with a high cost to movement. This approach, which has been used in many other studies, models landscape connectivity based on degree of naturalness or human modification and makes the assumption that more natural landscapes are less costly for many animals to move through and better facilitate ecological processes (Krosby et al., 2015; Spencer et al., 2010; Theobald et al., 2012). Pither et al. (2023) built their cost surface using the most up-to-date spatial data layers including the Canadian Human Footprint (Hirsh-Pearson et al., 2022) and an updated national road layer (Poley et al., 2022). We modified the 300-m cost surface of Pither et al. (2023) by adding a fifth lowest cost, which we assigned to natural areas within protected area boundaries under the assumption that these areas are less costly to move through than natural areas outside park boundaries (Spencer et al., 2010; Table 1). We reasoned that regulation of various human activities, such as resource extraction, hunting, and recreation, within protected areas as well as traditional land use practices and stewardship within Indigenous Protected and Conserved Areas (IPCAs) can provide benefits through decreased disturbance and mortality, allowing wildlife to move more easily through protected and conserved areas (Fryxell et al., 2020; Hebblewhite & Whittington, 2020; Indigenous Circle of Experts, 2018; Obbard et al., 2017; Schuster et al., 2019). With protected areas having the lowest cost on the landscape, adding a new park in the future, even if it is not a network node, will reduce the overall mean pairwise effective resistance relative to the network prior to addition of the park. We considered this to be an important property of an indicator for a protected areas network.

**Table 1.** List of cost values used to classify the cost surface and types of landscape features assigned each cost rank. See Pither et al. (2023) for a detailed description of landscape feature classifications and data layers used.

| cost value | landscape features |
| --- | --- |
| 0.1 | natural areas* within protected areas boundaries |
| 1 | natural areas* outside of protected areas boundaries |
| 10 | minor roads, pasturelands, forestry (cuts < 35 yrs old) |
| 100 | croplands, two-lane highways |
| 1000 | cities, railroads, multi-lane highways lakes ($\geq 10$ ha), rivers (flow > 28 m<sup>3</sup>/sec), mines, dams, nighttime lights |

\*natural areas refers to all terrestrial, non-anthropogenic landscape features including forests, wetlands, grasslands, etc.

The second input needed is a node file identifying the source and destination locations between which to evaluate connectivity. Often, source and destination nodes have been defined using a priori knowledge about core habitat patches that animals are most likely to move between and in the case of a park-to-park analysis, the parks serve as the nodes. More recently, omnidirectional methods have been developed which shift the nodes outside of the study area and are useful in cases where source and destination locations are unknown or for modelling landscape connectivity without reference to specific node locations (Koen et al., 2014; Phillips et al., 2021; Pither et al., 2023). Placing nodes within park boundaries is the most appropriate method for evaluating park-to-park connectivity; however, omnidirectional methods allow nodes to be independent of the parks network and therefore estimates of connectivity are repeatable over time. To evaluate connectivity of a protected areas network over time, we draw on properties of both methods to achieve our objective. We developed the sentinel node method, which involves randomly selecting a subset of parks to represent the full parks network. We use a single pixel placed within the park boundaries to represent a sentinel node. This allows an estimation of both internal park connectivity and connectivity between parks. Further, the consistent use of a randomly selected, fixed subset of park nodes allows changes in connectivity to be compared over time.

The following analyses were all performed using the Julia implementation of Circuitscape in pairwise mode (Julia version 1.7.3; Hall et al., 2021). Circuitscape provides two outputs which can be used to evaluate connectivity of a protected areas network. The first, a current density map, represents the probability of movement across the landscape and can be used to identify critical connectivity areas within or between protected areas. The second, pairwise effective resistance, is a measure of the cost of travelling between two pairs of park nodes for all pairs of nodes. The more potential pathways there are between nodes, the lower the effective resistance will be. We suggest the mean of the pairwise effective resistance (MPER) can serve as an indicator of overall network connectivity that can be tracked over time.

### 2.2. Representation of a full network and sensitivity to changes in the network

To determine how well the sentinel node method captures connectivity of the full park network, we used a small study area to model connectivity for a series of random park subsets and compared these to estimates of the full ‘true’ network from the same small area. We tested these methods in a ∼57,000 km^2^ area in eastern Ontario that represents 1 of 5 zones used to manage parks in the province. Using parks from the Canadian Protected and Conserved Areas Database (Environment and Climate Change Canada, 2023), which included provincial, federal, and private protected areas, we generated 3 random subsets of 20 parks in this small study area to represent 3 possible sentinel node scenarios. We then combined these 3 subsets, removing duplicates, to represent the full ‘true’ network, resulting in 52 parks. Park centroids were used as the location for nodes as opposed to the full park polygons in order to capture variability in connectivity within park boundaries (Brennan et al., 2022). For this analysis, all raster cells classified as natural cover (cost = 1) within park boundaries (both node parks and non-node parks) were re-assigned the lower cost of 0.1, representing the lowest cost on the landscape. All other cost values within park boundaries remained the same (10, 100, or 1000). Current density estimates of the 3 subsets and full network were compared using Spearman rank correlations of 1000 random pixel values.

We tested how well the sentinel node method could detect future changes in the network using 4 simulated scenarios in the small study area – two development scenarios that would reduce connectivity and two park-addition scenarios that would enhance connectivity. For the development scenarios, we first added a 157-km^2^ high-cost development (cost = 1000) in an area of relatively low cost with a high-cost road feature connecting it to an existing highway. In the second scenario, we added a 199-km^2^ development, in addition to the previous development. For the park-addition scenarios we (1) added a new 33-km^2^ protected area (cost = 0.1 for natural areas) to a heavily developed region on the landscape and (2) added a second 336-km^2^ protected area in addition to the first (cost = 0.1 for natural areas). We ran each of the 4 described scenarios for the 3 random sentinel node networks and the full ‘true’ network. To measure potential changes in connectivity with each of the new development scenarios, we calculated mean pairwise effective resistance (MPER) for all scenarios with each park network. An increase in MPER indicates a loss of connectivity across the network, while a decrease in MPER indicates improved connectivity.

### 2.3. Number of sentinel nodes required

Koen et al. (2014) found that for modelling omnidirectional connectivity, which uses source and destination nodes around the perimeter of a study area, a minimum of 20 nodes was required to accurately estimate current density throughout the study area using pairwise Circuitscape analysis. The authors showed that above this threshold, correlations between current density estimates and mean current density across the landscape reaches an asymptote, indicating additional nodes do not improve the connectivity model. We followed a similar approach to determine how many sentinel nodes are required to accurately represent the full park network.

To conduct this analysis, we used a ∼4,200 km^2^ study area containing ∼100,000 pixels. Using the 300-m cost surface of Pither et al. (2023), we randomly selected 60 locations in the study area and assigned points to these locations which we assumed to represent the true protected areas network. We then added 300-m buffers to these 60 spatial points, rasterized the buffered areas to a 300-m resolution, and reclassified as the lowest cost (0.1) to represent our full park network. We used simulated parks here to reduce any potential influence of park size and shape on the estimates of current density. Following the methods of Koen et al. (2014), we connected randomly selected combinations of park nodes starting with 2 pairs and sequentially increasing until we reached the full 60 node complement, resulting in 59 different cumulative current density maps. We measured mean current density for all maps as well as the correlation between values of each map (2 – 59 nodes) with values of the full network (60 nodes). A high correlation between values of a given map and the full map suggests that the spatial configuration of current density estimates is similar (Koen et al., 2014). In addition, we calculated and compared mean pairwise effective resistance for each iteration.

### 2.4. Case study: Assessing the current state of connectivity for Ontario’s protected areas network

We tested the sentinel node method on a larger landscape using the entire province of Ontario as a study area. Ontario covers an area of ∼1.07 million km^2^ and contains >1400 protected areas including provincial parks, federal parks, and private protected areas. Using a sentinel node analysis would not only help to reduce computational demands, but would also provide a baseline evaluation of the state of connectivity for the Ontario protected areas network and help inform future protected areas planning and design. For our provincial connectivity analysis, we used the 300-m resolution cost surface produced by Pither et al. (2023) cropped to the extent of Ontario, plus a ∼325-km wide buffer to reduce edge effects (Koen et al., 2010, 2014). For this province-wide analysis we assigned the lowest cost (0.1) to natural areas within parks and kept all other underlying landcover costs the same. Here, we make the assumption that natural lands within parks are less costly for animals to move through than natural lands outside of parks, while other landcover features within parks, such as roads, lakes, and built-up areas, remain costly.

Using our node analysis, we determined that 50 sentinel nodes would provide an accurate estimate of connectivity for the full park network. To initially select nodes, we limited the choice to only parks classified as ‘Provincial Parks’, since the analysis occurs within provincial boundaries and the province has jurisdiction over these parks; however, we integrated all classifications of protected areas (∼1423 PAs excluding protected portions of waterbodies) within the province into the cost surface. We also excluded parks with an area smaller than the size of a pixel (i.e., 300 m x 300 m) and parks located on islands from selection as a node. This resulted in a pool of 314 provincial parks from which to select 50 sentinel node parks. To ensure an even distribution of nodes across the province, we divided the province into three zones: northeast, northwest, and south (Fig. 2). To select the 50 park nodes, we used a stratified, random selection procedure with a set minimum proximity of 100 km between park centroids. Additionally, we specified that of the 50 parks, 20 each should be drawn from both the northeast and northwest zones and 10 from the south zone, which is roughly in proportion to the size of each zone. This selection procedure was implemented using the package *spatialEco* (v1.3-6; Evans & Murphy, 2021) in R (v4.2.2;R Core Team, 2022). Due to the number and proximity of parks within each zone, the selection algorithm converged on 45 parks representing 19, 14, and 12 parks within the northwest, northeast, and south zones, respectively. The remaining 5 parks were selected manually while ensuring sufficient distance from other parks. We used park centroids as node locations, provided centroids fell within a low cost (0.1) pixel. If the centroid fell within a higher cost pixel (1 – 1000), we adjusted the node location to a directly adjacent or the nearest, lowest cost (0.1) pixel. If a park had no pixels with a cost of 0.1, the node was placed in a pixel with the next available, lowest cost. Similar methods for node placement were used by Barnett and Belote (2021).

To map current density, which is an estimate of the probability of animal movement, we used pairwise mode Circuitscape in Julia with our input cost surface and 50 sentinel park nodes. Pairwise Circuitscape generates a cumulative current density map as an output by connecting and passing current between all possible pairs of nodes (1,225 pairs for 50 nodes). The resulting current density map can be used to identify important animal movement pathways between protected areas. We sampled 1000 random locations to compare our estimate of current density across Ontario to a recent estimate made by Pither et al. (2023) at the same extent and resolution. Circuitscape also calculates pairwise effective resistance between all pairs of nodes, which is a measure of cumulative resistance between connected nodes. The more potential pathways between nodes, the lower the pairwise effective resistance and thus, greater connectivity. We calculated the mean pairwise effective resistance (MPER) across the 50 sentinel nodes as a measure of overall network connectivity.

### 2.5. Node isolation

To measure isolation of individual node parks, we calculated the mean pairwise effective resistance between each sentinel node and all other nodes. This node isolation metric is similar to the PAI index used by Brennan et al. (2022) to measure the degree of isolation of global protected areas. Mean node isolation values can help identify parks within the network that are least connected and thus highlights areas where conservation interventions are most likely to help improve overall network connectivity.

## 3. Results

### 3.1. Representation of a full network and sensitivity to changes in the network

Within the small study area in eastern Ontario, current density values from maps produced with randomly selected subsets of park nodes were highly correlated with each other and with current density values of a full “true” park network (mean *rho* = 0.91, range = 0.83 – 0.97). Mean pairwise effective resistance (MPER) between sentinel nodes, as a connectivity indicator, was able to detect simulated land-use and landcover changes. Across all random scenarios and the true scenario, MPER increased with the addition of a high-cost development and increased further with addition of a second high-cost development (Table 2). MPER decreased, indicating enhanced connectivity, with the addition of a new, low-cost protected area across all scenarios and decreased more sharply with the addition of a second protected area (Table 2). In neither case was the new park a node in the analysis, so the benefit to connectivity was achieved through the addition of new lower cost areas to the landscape.

**Table 2.** Mean pairwise effective resistance values (ohms) of 3 random sentinel node scenarios (20 nodes/scenario; R1, R2, and R3) and the full “true” park network (52 nodes) for (1) the original cost surface; (2) addition of a high-cost human development; (3) addition of a second high-cost development; (4) the addition of a new low-cost protected area; and (5) addition of a second protected area. Effective resistance is a measure of the cumulative cost of moving between a pair of nodes. Mean pairwise effective resistance is an indicator of overall network connectivity.

| scenario | no<br>development | single<br>development | multiple<br>developments | single<br>park | multiple<br>parks |
| --- | --- | --- | --- | --- | --- |
| R1 | 116.102 | 116.164 | 116.623 | 116.101 | 116.078 |
| R2 | 139.772 | 139.809 | 140.401 | 139.767 | 139.751 |
| R3 | 280.496 | 280.528 | 280.960 | 280.490 | 280.474 |
| Full | 181.642 | 181.685 | 182.154 | 181.638 | 181.620 |

### 3.2. Number of sentinel nodes required

We determined that 50 sentinel nodes was sufficient to accurately depict current density estimates of a simulated park network. Spearman rank correlations between current density values from maps with increasing numbers of nodes compared to values from the full map (60 nodes) reached an asymptote at around 50 nodes (Fig 1a; *rho* = 0.99 for the comparison between 50 and 60 nodes). Similarly, variation in mean pairwise effective resistance values was minimized at about 50 nodes (Fig 1b).

**Fig 1.**
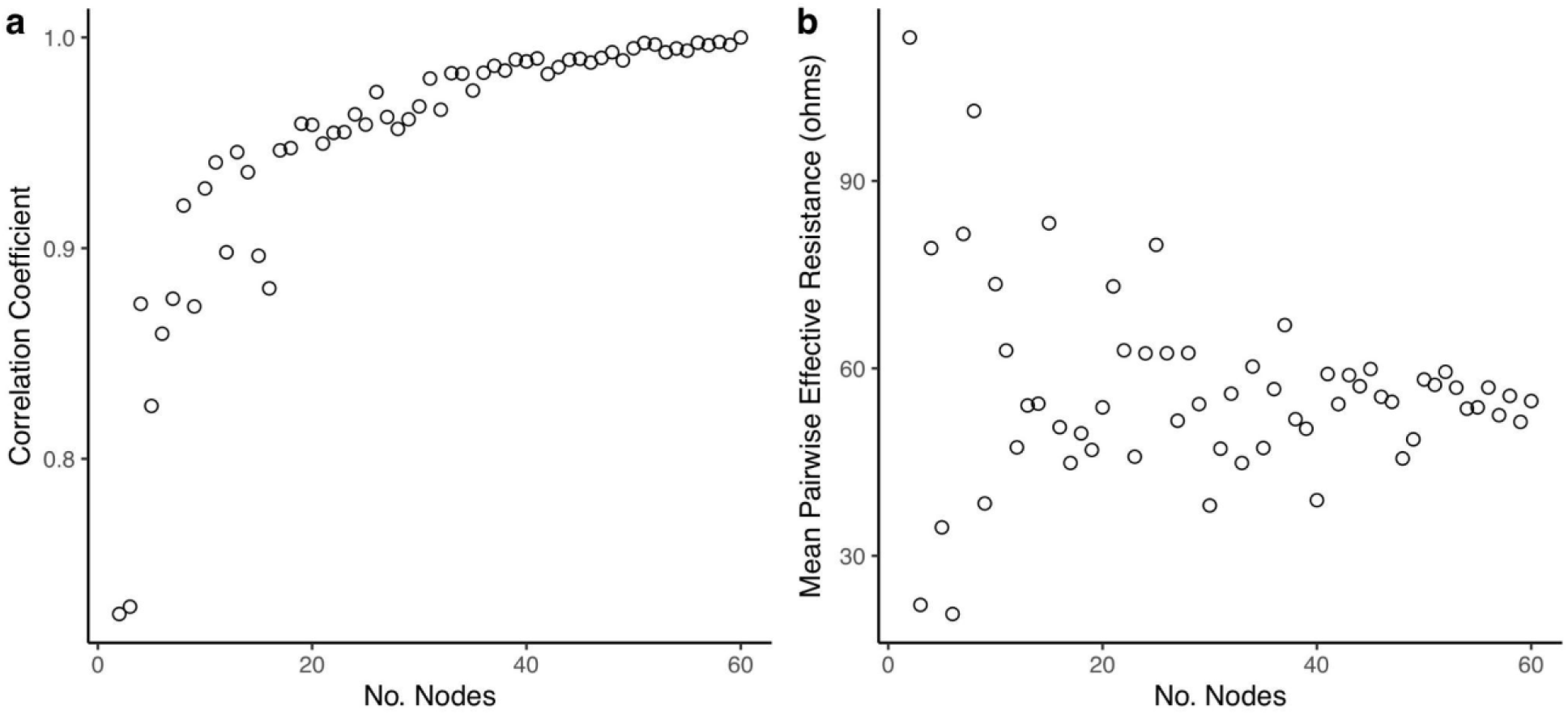
a) Spearman rank correlation coefficients of current density maps using increasing numbers of nodes compared to a current density map using all possible node connections. Number of nodes connected ranges from 2 – 60. All correlations are based on 1000 randomly drawn current density values; b) Mean pairwise effective resistance values calculated for sentinel park node networks with increasing numbers of node pairs (range = 2 – 60 nodes).

### 3.3. Case study: Assessing the current state of connectivity for Ontario’s protected areas network

The final set of 50 sentinel nodes contained 20, 15, and 15 parks within the northwest, northeast, and south zones of Ontario, respectively (Fig 2). The cumulative current density map for the full province displays varied patterns of current density between protected areas across the province (Fig 3). For example, many areas of high current density (i.e., pinch points) can be seen between parks connecting the south of the province to the north (Fig 4b). This tendency for a pinching of current between south and north is magnified by the Great Lakes, as all current needs to pass to the north of the lakes. A lack of park-to-park connectivity is evident in the south of the province (Fig 4c) where small protected areas are isolated by a matrix of human development. In contrast, areas of low, diffuse current density, where many potential movement pathways exist, can be found in the intact, low-cost northern regions of the province (Fig 4a). There was a positive correlation (Spearman’s *rho* = 0.76, p < 0.001) between values from our park-to-park current density map and values from the omnidirectional current density map produced by Pither et al. (2023).

**Fig 2.**
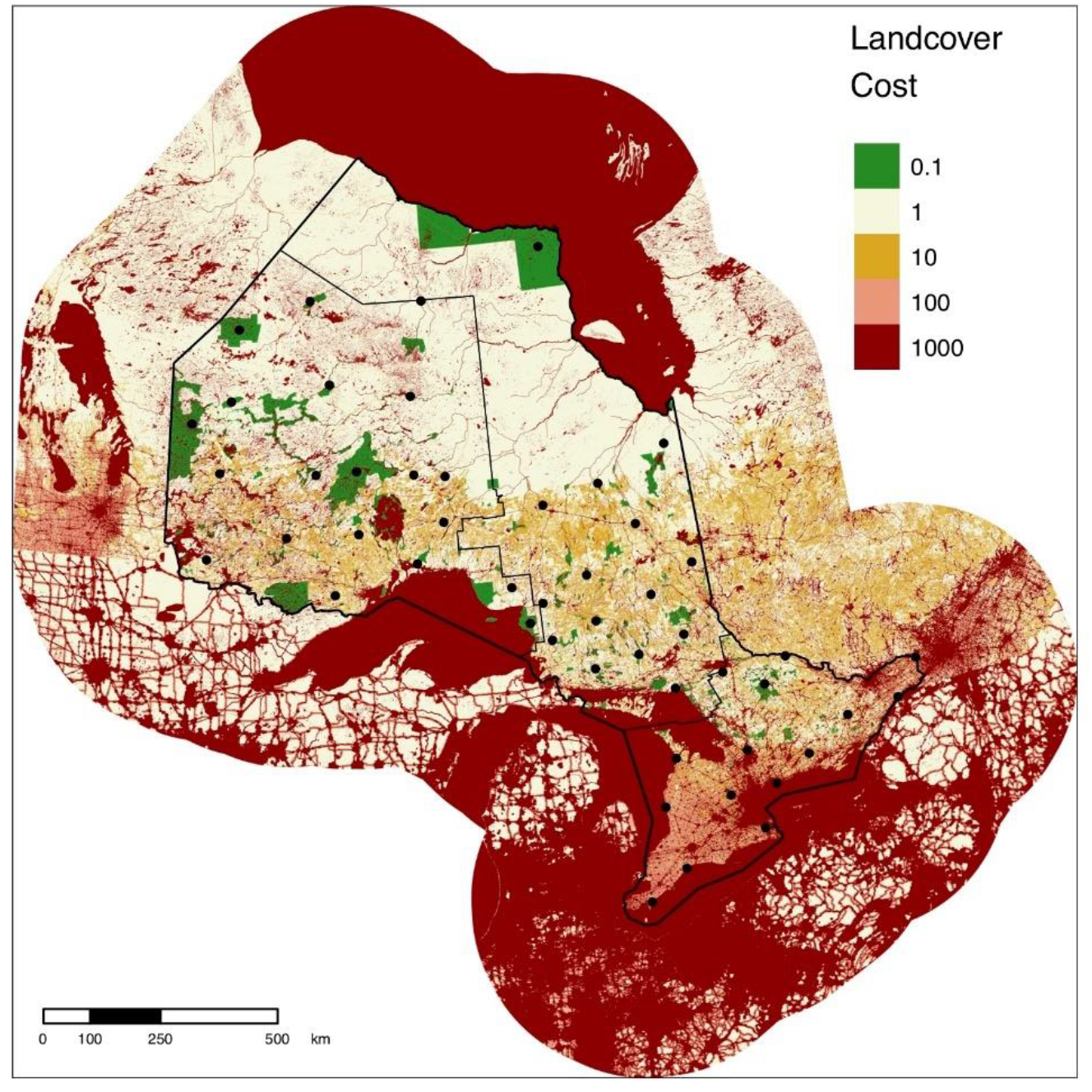
Cost surface for Ontario with sentinel nodes. Five cost values were assigned to landscape features based on the degree to which they facilitate or impede movement. Natural areas within protected areas boundaries were assigned the lowest cost (0.1) under the assumption that they are less costly to move through than natural areas outside of protected areas. Solid black circles show the locations of the 50 sentinel nodes used to evaluate protected areas connectivity in Ontario. Solid black lines show the boundaries of the zones used to stratify the province for node selection. The modified cost surface was derived from the 300-m resolution cost surface of Canada from Pither et al. (2023).

**Fig 3.**
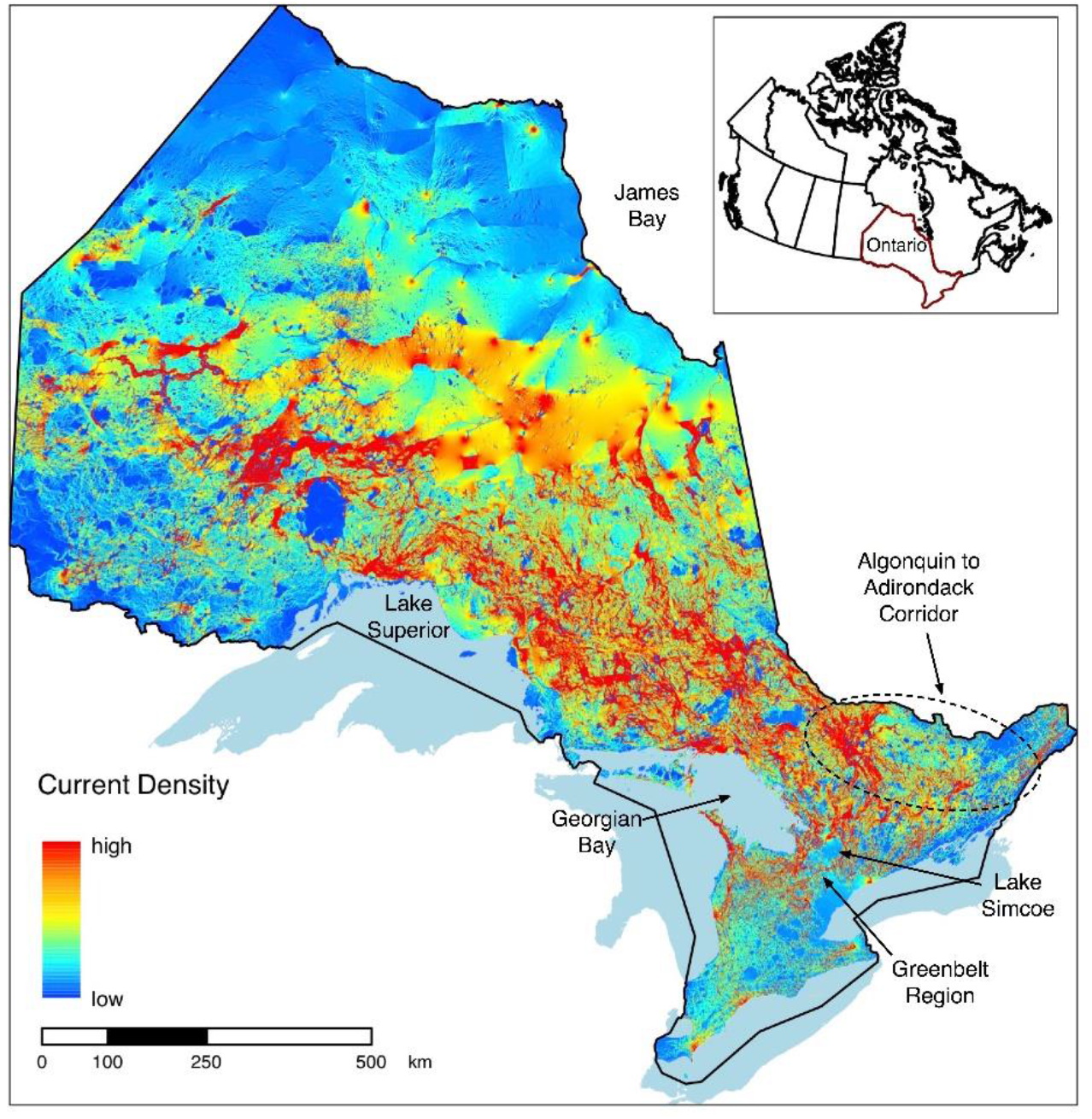
Current density map displaying park-to-park connectivity for the protected areas network in Ontario. Current density, measured in amps (A), represents the probability of animal movement within a given pixel across the landscape. The dashed oval shows the general extent of the Algonquin to Adirondack Corridor.

**Fig 4.**
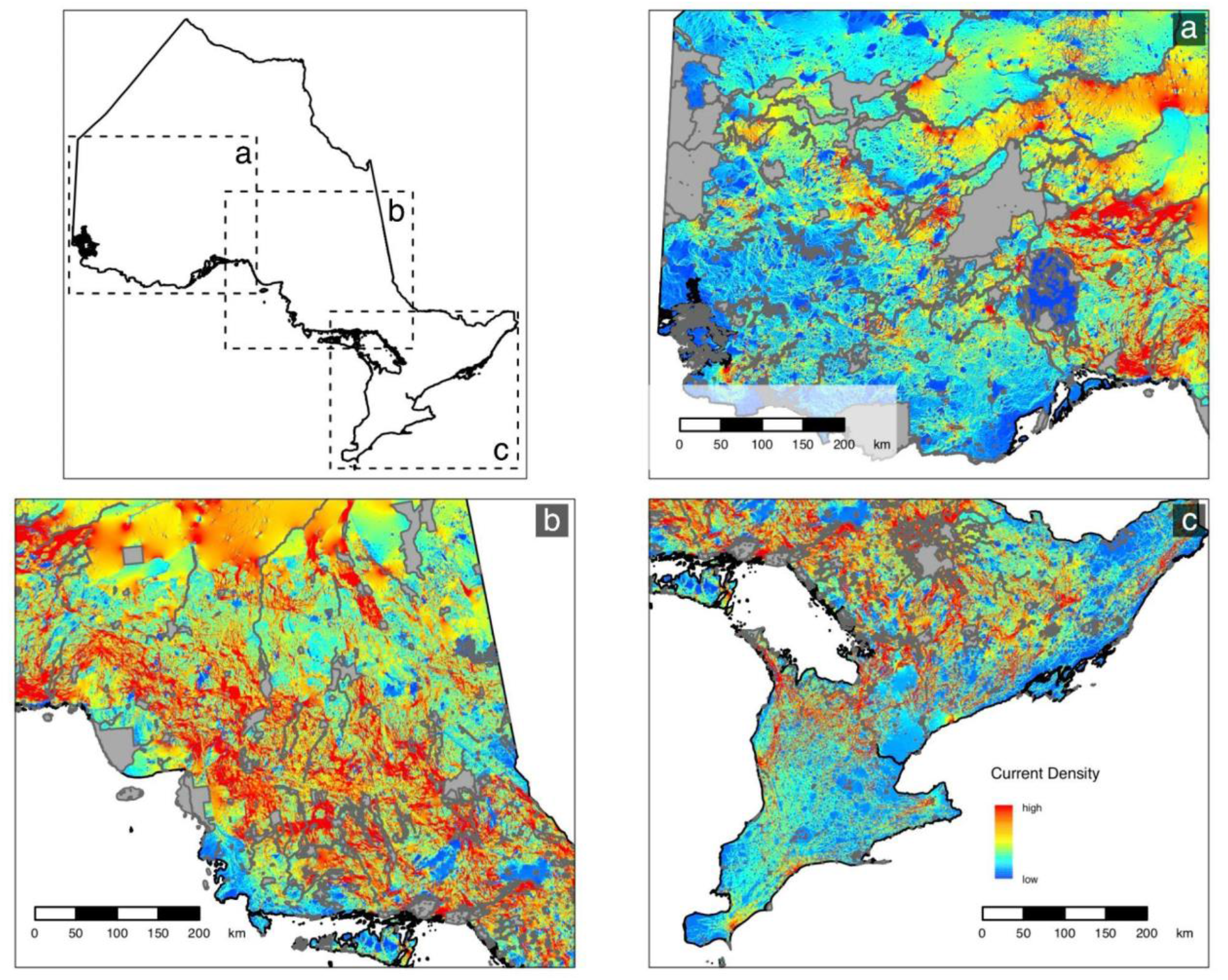
Vignettes showing three areas that highlight differences in protected areas connectivity across the provinces. Regions displayed are (a) northwestern Ontario (b) central Ontario; and (c) southern Ontario. Grey polygons show protected and conserved areas.

Mean (SE) pairwise effective resistance among the sentinel nodes was 83.07 (3.07) ohms. Pairwise effective resistance (also termed resistance distance) and Euclidean distances varied among pairs of nodes and there was a weak positive relationship between Euclidean distance and resistance distance (*rho* = 0.34, p < 0.001; Fig 5). Node isolation values were positively skewed (Fig 6a) and the highest isolation values were found in the south of the province (Fig 6b; Table S1).

**Fig 5.**
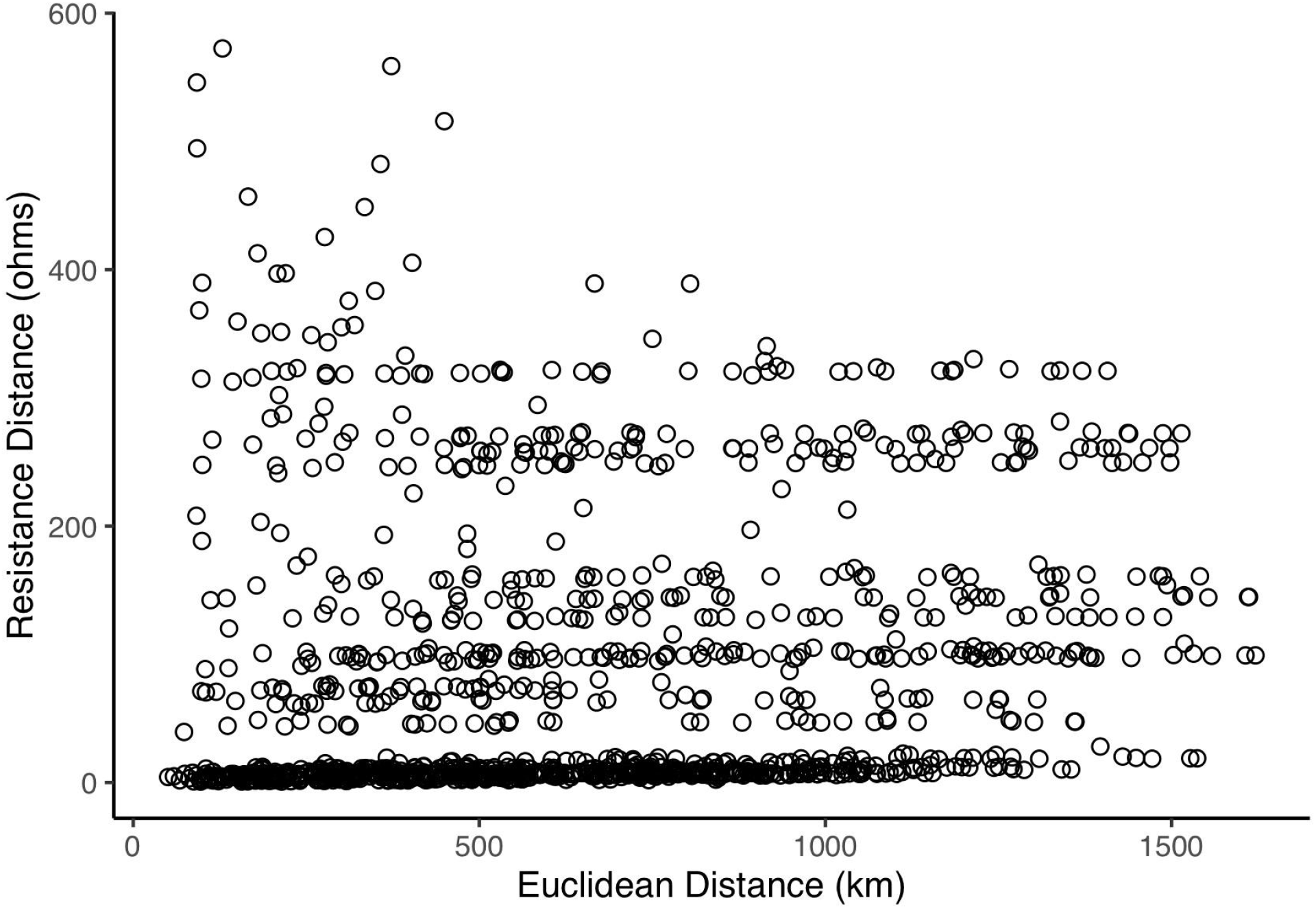
Relationship between pairwise Euclidean distance and resistance distance of sentinel nodes.

**Fig 6.**
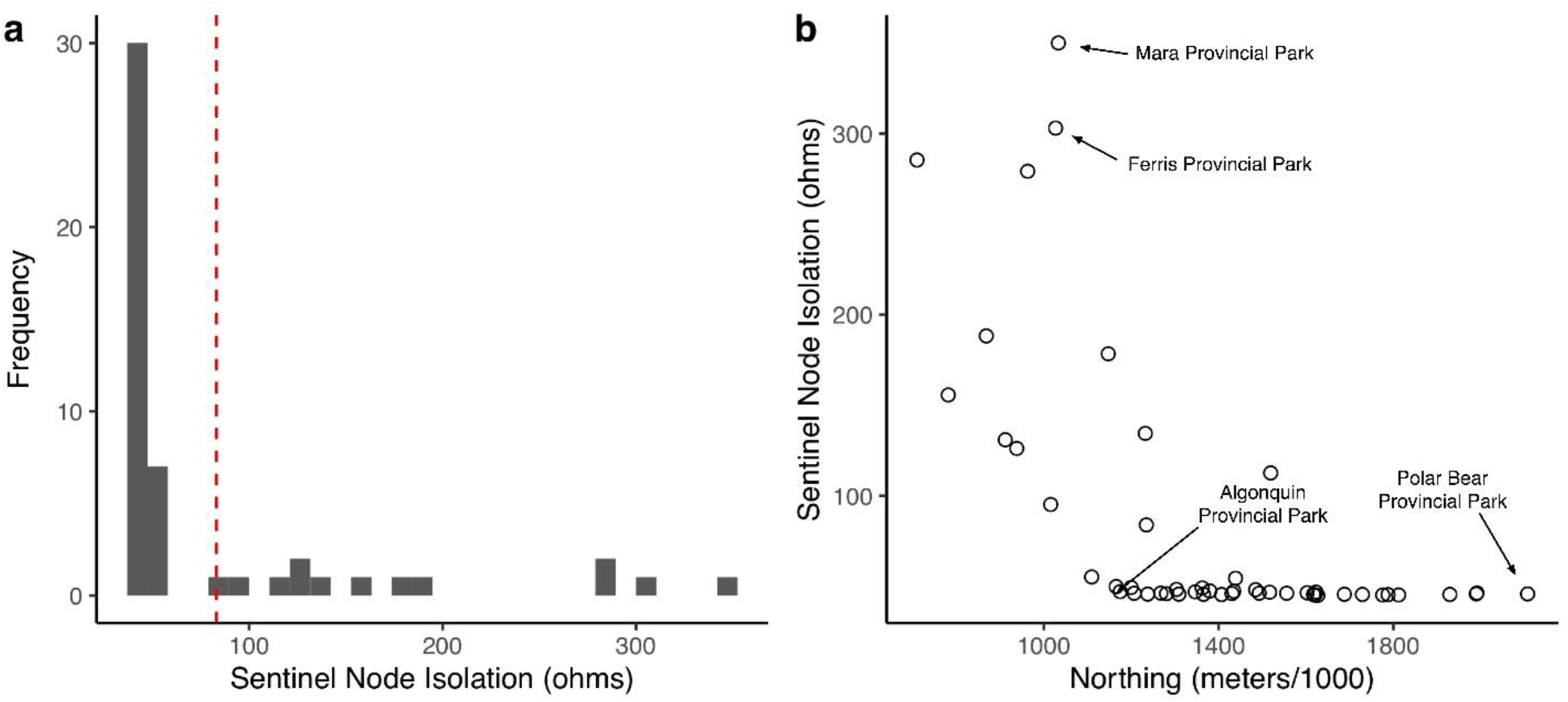
a) Frequency distribution of sentinel node isolation values; higher values indicate a higher degree of isolation. The red dashed line shows the mean pairwise effective resistance for all 50 sentinel nodes (MPER = 83.07 ohms). b) Relationship between sentinel node isolation and northing values.

## 4. Discussion

We developed a new method using circuit theory to evaluate connectivity of a protected areas network, and to establish an indicator of changing connectivity over time. In contrast to many other park-to-park connectivity models, we assigned a lower cost to protected areas in our cost surface and we use a fixed subset of protected areas as sentinel nodes to represent the full protected areas network. Our method was sensitive to added developments, where the indicator increased as expected, and added parks, where the indicator decreased. Our method can be used to assess the current state of connectivity for a given protected areas network and track changes in connectivity through time. As nations push to protect 30% of terrestrial lands by 2030 in well-connected protected areas networks, the sentinel node method of measuring park-to-park connectivity provides a consistent and repeatable framework to incorporate connectivity into conservation strategies, and in particular, protected areas planning.

Our findings suggest that the sentinel node method is able to detect land cover and land-use changes. Specifically, we found that MPER was effective as a connectivity indicator to track changes in a protected areas network. Under different simulated scenarios, MPER increased in response to added high-cost developments and decreased in response to addition of new protected areas in comparison to current estimates of connectivity. We consider that MPER meets many of the desirable properties of a connectivity indicator outlined by Theobald et al. (2022). In particular, MPER incorporates interpatch distance (i.e., resistance distance), reflects both within-and between-patch connectivity, can be calculated at a high resolution (<1km) with relative computational efficiency, and is simple to interpret. We plan to conduct more simulations to test the sensitivity of the sentinel node method, which would help in determining how well MPER, as a connectivity indicator, fits other properties outlined by Theobald et al. (2022). We believe MPER and our measure of node isolation are valuable additions to other indicators of functional connectivity (e.g., PAI index; Brennan et al., 2022) and in combination with a second type of output from our analyses, current density maps that highlight areas with a high probability of animal movement, can help practitioners achieve progress towards conservation targets.

Theobald et al. (2022) highlight that a connectivity indicator should be computationally efficient and simple to re-estimate in the future. Circuit theory models have become increasingly popular for evaluating connectivity; however, with these resistance surface-based connectivity metrics, computational demand increases as the number of pixels in a cost surface increases and with an increasing number of nodes. Thus, it may become unrealistic for protected areas ecologists and planners to evaluate park-to-park connectivity at a high resolution (<1km) over a regional or national extent for an entire protected areas network. Koen et al. (2014) showed that 20-30 nodes were needed around the perimeter of their study area to produce accurate estimates of connectivity using omnidirectional methods. They suggested similar sensitivity analyses should be undertaken by other researchers to ensure sufficient sampling for their study.

Following these recommendations, we determined that 50 nodes should be sufficient to accurately represent a full protected areas network when assessing park-to-park connectivity. Researchers may wish to test the generality of this finding in networks of varying structure. We suggest that using a subset of sentinel nodes reduces the computational demand of running circuit theory models, allowing ecologists and planners to evaluate protected areas connectivity more efficiently while also providing a static network with which to track future connectivity.

We used our sentinel node method to evaluate connectivity of the protected areas network for the province of Ontario. The cumulative current density map shows differing patterns of current density among protected areas across the province. As expected, connectivity is lower among protected areas in the south of the province, where small parks are isolated within a matrix of urban development and agriculture. Protection and restoration efforts (e.g., tree-planting and grassland restoration) have a high potential to maintain and improve connectivity within this region. Despite the high anthropogenic impact in the south, a number of high current density areas can be seen flowing south to north between Georgian Bay and Lake Simcoe, along Ontario’s Greenbelt region, and west to east along the Algonquin to Adirondack corridor. Continued expansion of urban development makes high connectivity areas in this region particularly vulnerable to loss and thus priorities for protection. Many sites have been identified as candidate Key Biodiversity Areas (KBAs) in the south of the province (http://www.kbacanada.org). These KBAs could serve as important candidates for new protected areas that would make significant contributions to conserving biodiversity and maintaining connectivity.

A concentration of high current density is evident in the region connecting the south of the province to the north. Many of these areas align with previous omnidirectional models of connectivity (Bowman & Cordes, 2015; Pither et al., 2023) including high current flow north of Lake Superior and a corridor moving east to west from James Bay towards the Ontario-Manitoba border. The high concentration of current flow in this region is partly an artifact of park-to-park connectivity models, however the spatial configuration of high current can be attributed to the geometry and geography of the province, and to landscape heterogeneity. This part of the province contains a mix of natural boreal forests, lake and rivers, urban development, and natural resource extraction (e.g., commercial forestry and mining). The close proximity of low and high cost land cover features leads to the funneling of current through this region as evidenced by many high current pinch points (Marrotte et al., 2017). While there are many similarities between our park-to-park model of connectivity and omnidirectional models, notable differences can be seen in the high current flowing through protected areas as a result of the low cost assigned to natural areas within park boundaries. As species distributions shift in response to climate change, these pinch points connecting the south of the province to the north will be critical for maintaining climate connectivity (Parks et al., 2023; Schloss et al., 2022) and encroachment of surrounding anthropogenic developments puts these critical connectivity areas at risk of being reduced or lost. Connectivity in this region could be maintained through a combination of expanded protection and improved management of working landscapes between protected areas.

In contrast to the south, protected areas in the northeast and northwest regions of the province are surrounded by a generally intact, natural landscape. Low to moderate current flow in these areas does not indicate low connectivity, but rather that many potential movement pathways exist and therefore current flow is more diffuse. While currently less vulnerable to anthropogenic threats, these areas are as important to protect in order to prevent fragmentation of these remaining intact landscapes, especially since protection of intact landscapes can provide more immediate and cost effective conservation benefits than restoration of degraded habitats (Cook-Patton et al., 2021). Further, the province’s north contains some of the largest remaining global peatlands, which store vast amounts of irrecoverable carbon (Mitchell et al., 2021; Sothe et al., 2022) and if lost would result in significant carbon debt (Goldstein et al., 2020; Noon et al., 2021). O’Brien et al. (2023) mapped the overlap between carbon stocks and omnidirectional current density estimates in Ontario to identify areas important for nature-based climate solutions. Similar methods could be employed using the results from the current analysis to identify hotspots specific to protected areas connectivity to help guide identification of new candidate PAs. Five IPCAs have been proposed by First Nations across Ontario, three of which occur in the northwest and northeast region of the province (Environment and Climate Change Canada, 2021). Establishment of these protected areas would contribute to connectivity of the protected areas network, provide benefits towards climate change mitigation, and help to uphold Indigenous rights by shifting away from traditional, colonial concepts of conservation and towards those led by Indigenous communities and that integrate Indigenous culture and priorities. Continued collaboration and support of Indigenous-led conservation efforts to secure traditionally used, governed, and conserved lands and waters should remain a high priority as we strive to reach ambitious biodiversity and climate targets (Moola & Roth, 2019).

In addition to producing a current density map, which helps to identify areas important for connectivity, we also calculated MPER as an indicator of connectivity for the Ontario protected areas network. As this is the first time protected areas connectivity has been evaluated in the province, this serves as a baseline evaluation of connectivity for which future estimates can be compared to track changes in connectivity through time. Brennan et al. (2022) found that a combination of restoration and protected areas expansion maximized reductions in their protected areas isolation (PAI) index, thus improving connectivity. Similarly, we found that addition of protected areas could decrease MPER of our parks network. Our results and previous work suggests that land-use planners and conservation practitioners could maintain and enhance connectivity of protected areas networks through a combination of protected areas expansion, improved land management practices, and restoration efforts. Expanding current protected areas and enhancing connectivity between protected areas can help make significant contributions to global targets to conserve biodiversity (Target 3; Convention on Biological Diversity, 2022).

This will be especially critical in the face of climate change as currently suitable habitat in protected areas may become unsuitable in the future and species are required to shift their ranges to track suitable climates (Parks et al., 2023). We suggest that future work should more rigorously test the sentinel node method by examining the response of MPER to protected areas size and configuration and different conservation interventions (e.g., addition of corridors).

We note that a few issues arise when attempting to re-evaluate a connectivity indicator for a protected areas network over time that are important to recognize. For example, a protected areas network is, and should be, dynamic over time as new protected areas are added, existing areas expanded, or new management practices implemented. These changes in the network across space and time impact the consistency of future estimates; however, the use of a static, unchanging set of sentinel nodes as we present here provides a repeatable framework to measure connectivity of an ever-changing network. Second, future changes in the network can become conflated with changes in the availability of new land cover data and technology or methods used to collect these data. To reduce the likelihood of these issues affecting future assessments of connectivity, we suggest that (1) cost surface parameterization should follow the same cost ranking scheme (see Pither et al., 2023); and (2) the resolution of future analyses should match that of previous analyses, which may require aggregating new, higher resolution data to coarser resolutions (in this case 300 m).

The mean pairwise effective resistance, as an indicator of connectivity, was found to be 83.07 ± 3.07 (mean ± se) for the Ontario protected areas network. This is a baseline estimate of the proposed connectivity indicator, which will be informative to compare over time to future assessments of the network. For now, we can use our analysis to identify that there is spatial variation across the province in the degree of connectivity, and suggest that to enhance connectivity of the protected areas network, a target would be to reduce MPER over time (i.e., shift the estimate towards the lower range of cost values). This can be accomplished by reducing costs on the map in a variety of ways. Perhaps the most efficient way to reduce MPER would be to add protected areas in or near anthropogenically developed landscapes. Evaluating alternative strategies for reducing MPER through simulation will undoubtedly be a useful strategy for landscape managers. With national targets to increase protection of terrestrial lands to 25% by 2025 and 30% by 2030, we suggest that it would be reasonable for connectivity of the Ontario protected areas network to be re-evaluated in conjunction with these targets to track progress.

In addition to the mean pairwise effective resistance, we also calculated isolation values for individual sentinel nodes, which is comparable to the Protected Areas Isolation (PAI) index used by Brennan et al. (2022). Our node isolation index confirms that there is spatial variation in connectivity across the province with the highest degree of node isolation occurring in the south. Further, estimates of node isolation help to distinguish between nodes occurring in relatively intact, natural landscapes and those in heavily human modified landscapes. Both areas can have low current density values, but the latter will have a high isolation value (i.e., high degree of isolation), while the former will have a low isolation value owing to many potential movement pathways throughout intact natural landscapes. We suggest that this index of node isolation could be used to identify individual nodes where restoration work or other landscape management within and around a given protected area would help to improve overall network connectivity.

The recognized importance of connectivity for biodiversity is evident from the adoption of a new ambitious global biodiversity agreement including increased targets to protect 30% of terrestrial lands in ecologically representative and well-connected networks of protected areas. Key to achieving this target is the ability to measure connectivity and to track progress. We present a method through which government and non-government organizations can evaluate connectivity of protected areas networks over time within a repeatable framework. This will allow connectivity to be better incorporated into protected areas planning, which is vital to creating functional networks of protected areas.

## Acknowledgements

Financial support for this study was provided by the Ontario Ministry of Environment, Conservation, and Parks for P. O. and N.C., and by the Ontario Ministry of Natural Resources and Forestry for J.B. Thank you to additional members of the Ontario Parks connectivity working group Karen Hartley, Louis Chora and Amanda Schroeder for advice. We also thank Richard Pither, Angela Brennan, and Kristen Hirsh-Pearson for collaboration on related work and Jochen Jaeger for the insightful discussion.

## Data Availability

Our input movement cost surface layer and sentinel node location file and output current density map are available at Figshare repository doi: XXXX.

## Literature Cited

Andam, K. S., Ferraro, P. J., Pfaff, A., Sanchez-Azofeifa, G. A., & Robalino, J. A. (2008). Measuring the effectiveness of protected area networks in reducing deforestation. Proceedings of the National Academy of Sciences, 105(42), 16089–16094. https://doi.org/10.1073/pnas.0800437105

Barnett, K., & Belote, R. T. (2021). Modeling an aspirational connected network of protected areas across North America. Ecological Applications, 31(6). https://doi.org/10.1002/eap.2387

Barrueto, M., Forshner, A., Whittington, J., Clevenger, A. P., & Musiani, M. (2022). Protection status, human disturbance, snow cover and trapping drive density of a declining wolverine population in the Canadian Rocky Mountains. Scientific Reports, 12(1), 17412. https://doi.org/10.1038/s41598-022-21499-4

Bellard, C., Bertelsmeier, C., Leadley, P., Thuiller, W., & Courchamp, F. (2012). Impacts of climate change on the future of biodiversity: Biodiversity and climate change. Ecology Letters, 15(4), 365–377. https://doi.org/10.1111/j.1461-0248.2011.01736.x

Belote, R. T., Dietz, M. S., McRae, B. H., Theobald, D. M., McClure, M. L., Irwin, G. H., McKinley, P. S., Gage, J. A., & Aplet, G. H. (2016). Identifying Corridors among Large Protected Areas in the United States. PLOS ONE, 11(4), e0154223. https://doi.org/10.1371/journal.pone.0154223

Bowman, J., Adey, E., Angoh, S. Y. J., Baici, J. E., Brown, M. G. C., Cordes, C., Dupuis, A. E., Newar, S. L., Scott, L. M., & Solmundson, K. (2020). Effects of cost surface uncertainty on current density estimates from circuit theory. PeerJ, 8, e9617. https://doi.org/10.7717/peerj.9617

Bowman, J., & Cordes, C. (2015). Landscape Connectivity in the Great Lakes Basin. figshare. https://doi.org/10.6084/M9.FIGSHARE.1471658.V1

Brennan, A., Naidoo, R., Greenstreet, L., Mehrabi, Z., Ramankutty, N., & Kremen, C. (2022). Functional connectivity of the world’s protected areas. Science, 376(6597), 1101–1104. https://doi.org/10.1126/science.abl8974

Butchart, S. H. M., Scharlemann, J. P. W., Evans, M. I., Quader, S., Aricò, S., Arinaitwe, J., Balman, M., Bennun, L. A., Bertzky, B., Besançon, C., Boucher, T. M., Brooks, T. M., Burfield, I. J., Burgess, N. D., Chan, S., Clay, R. P., Crosby, M. J., Davidson, N. C., De Silva, N., … Woodley, S. (2012). Protecting Important Sites for Biodiversity Contributes to Meeting Global Conservation Targets. PLoS ONE, 7(3), e32529. https://doi.org/10.1371/journal.pone.0032529

Carroll, C., & Ray, J. C. (2021). Maximizing the effectiveness of national commitments to protected area expansion for conserving biodiversity and ecosystem carbon under climate change. Global Change Biology, 27(15), 3395–3414. https://doi.org/10.1111/gcb.15645

Chen, I.-C., Hill, J. K., Ohlemüller, R., Roy, D. B., & Thomas, C. D. (2011). Rapid Range Shifts of Species Associated with High Levels of Climate Warming. Science, 333(6045), 1024– 1026. https://doi.org/10.1126/science.1206432

Convention on Biological Diversity. (2022). Kunming-Montreal Global biodiversity framework. https://www.cbd.int/doc/c/e6d3/cd1d/daf663719a03902a9b116c34/cop-15-l-25-en.pdf

Cook-Patton, S. C., Drever, C. R., Griscom, B. W., Hamrick, K., Hardman, H., Kroeger, T., Pacheco, P., Raghav, S., Stevenson, M., Webb, C., Yeo, S., & Ellis, P. W. (2021). Protect, manage and then restore lands for climate mitigation. Nature Climate Change, 11(12), 1027-1034–0. https://doi.org/10.1038/s41558-021-01198-0

Craigie, I. D., Baillie, J. E. M., Balmford, A., Carbone, C., Collen, B., Green, R. E., & Hutton, J. M. (2010). Large mammal population declines in Africa’s protected areas. Biological Conservation, 143(9), 2221–2228. https://doi.org/10.1016/j.biocon.2010.06.007

Deslauriers, M. R., Asgary, A., Nazarnia, N., & Jaeger, J. A. G. (2018). Implementing the connectivity of natural areas in cities as an indicator in the City Biodiversity Index (CBI). Ecological Indicators, 94, 99–113. https://doi.org/10.1016/j.ecolind.2017.02.028

Dickson, B. G., Albano, C. M., McRae, B. H., Anderson, J. J., Theobald, D. M., Zachmann, L. J., Sisk, T. D., & Dombeck, M. P. (2017). Informing Strategic Efforts to Expand and Connect Protected Areas Using a Model of Ecological Flow, with Application to the Western United States: Mapping ecological flow to inform planning. Conservation Letters, 10(5), 564–571. https://doi.org/10.1111/conl.12322

Doyle, P. G., & Snell, J. L. (1984). Random walks and electric networks. American Mathematical Society.

Environment and Climate Change Canada. (2021). Canada Target 1 Challenge. https://www.canada.ca/en/environment-climate-change/services/nature-legacy/canada-target-one-challenge.html#events

Environment and Climate Change Canada. (2023). Canadian Protected and Conserved Areas Database. https://www.canada.ca/en/environment-climate-change/services/national-wildlife-areas/protected-conserved-areas-database.html

Evans, J., & Murphy, M. (2021). _spatialEco_ [R package version 1. 3-6]. <https://github.com/jeffreyevans/spatialEco>

Fryxell, J. M., Avgar, T., Liu, B., Baker, J. A., Rodgers, A. R., Shuter, J., Thompson, I. D., Reid, D. E. B., Kittle, A. M., Mosser, A., Newmaster, S. G., Nudds, T. D., Street, G. M., Brown, G. S., & Patterson, B. (2020). Anthropogenic Disturbance and Population Viability of Woodland Caribou in Ontario. The Journal of Wildlife Management, 84(4), 636–650. https://doi.org/10.1002/jwmg.21829

Geldmann, J., Manica, A., Burgess, N. D., Coad, L., & Balmford, A. (2019). A global-level assessment of the effectiveness of protected areas at resisting anthropogenic pressures. Proceedings of the National Academy of Sciences, 116(46), 23209–23215. https://doi.org/10.1073/pnas.1908221116

Goldstein, A., Turner, W. R., Spawn, S. A., Anderson-Teixeira, K. J., Cook-Patton, S., Fargione, J., Gibbs, H. K., Griscom, B., Hewson, J. H., Howard, J. F., Ledezma, J. C., Page, S., Koh, L. P., Rockström, J., Sanderman, J., & Hole, D. G. (2020). Protecting irrecoverable carbon in Earth’s ecosystems. Nature Climate Change, 10(4), 287–295. https://doi.org/10.1038/s41558-020-0738-8

Goodwin, B. J., & Fahrig, L. (2002). How does landscape structure influence landscape connectivity? Oikos, 99(3), 552–570. https://doi.org/10.1034/j.1600-0706.2002.11824.x

Gray, C. L., Hill, S. L. L., Newbold, T., Hudson, L. N., Börger, L., Contu, S., Hoskins, A. J., Ferrier, S., Purvis, A., & Scharlemann, J. P. W. (2016). Local biodiversity is higher inside than outside terrestrial protected areas worldwide. Nature Communications, 7(1), 12306. https://doi.org/10.1038/ncomms12306

Haddad, N. M., Brudvig, L. A., Clobert, J., Davies, K. F., Gonzalez, A., Holt, R. D., Lovejoy, T. E., Sexton, J. O., Austin, M. P., Collins, C. D., Cook, W. M., Damschen, E. I., Ewers, R. M., Foster, B. L., Jenkins, C. N., King, A. J., Laurance, W. F., Levey, D. J., Margules, C. R., … Townshend, J. R. (2015). Habitat fragmentation and its lasting impact on Earth’s ecosystems. Science Advances, 1(2), e1500052. https://doi.org/10.1126/sciadv.1500052

Hall, K. R., Anantharaman, R., Landau, V. A., Clark, M., Dickson, B. G., Jones, A., Platt, J., Edelman, A., & Shah, V. B. (2021). Circuitscape in julia: Empowering dynamic approaches to connectivity assessment. Land, 10(3). https://doi.org/10.3390/land10030301

Hebblewhite, M., & Whittington, J. (2020). Wolves without borders: Transboundary survival of wolves in Banff National Park over three decades. Global Ecology and Conservation, 24, e01293. https://doi.org/10.1016/j.gecco.2020.e01293

Heller, N. E., & Zavaleta, E. S. (2009). Biodiversity management in the face of climate change: A review of 22 years of recommendations. Biological Conservation, 142(1), 14–32. https://doi.org/10.1016/j.biocon.2008.10.006

Hilborn, R., Arcese, P., Borner, M., Hando, J., Hopcraft, G., Loibooki, M., Mduma, S., & Sinclair, A. R. E. (2006). Effective Enforcement in a Conservation Area. Science, 314(5803), 1266–1266. https://doi.org/10.1126/science.1132780

Hilty, J., Worboys, G. L., Keeley, A., Woodley, S., Lausche, B. J., Locke, H., Carr, M., Pulsford, I., Pittock, J., White, J. W., Theobald, D. M., Levine, J., Reuling, M., Watson, J. E. M., Ament, R., & Tabor, G. M. (2020). Guidelines for conserving connectivity through ecological networks and corridors (C. Groves, Ed.). IUCN, International Union for Conservation of Nature. https://doi.org/10.2305/IUCN.CH.2020.PAG.30.en

Hirsh-Pearson, K., Johnson, C. J., Schuster, R., Wheate, R. D., & Venter, O. (2022). Canada’s human footprint reveals large intact areas juxtaposed against areas under immense anthropogenic pressure. Facets, 7, 398–419.

Hoffmann, A. A., Miller, A. D., & Weeks, A. R. (2021). Genetic mixing for population management: From genetic rescue to provenancing. Evolutionary Applications, 14(3), 634–652. https://doi.org/10.1111/eva.13154

Hooftman, D. A. P., Edwards, B., & Bullock, J. M. (2016). Reductions in connectivity and habitat quality drive local extinctions in a plant diversity hotspot. Ecography, 39(6), 583– 592. https://doi.org/10.1111/ecog.01503

Indigenous Circle of Experts. (2018). We Rise Together: Achieving Pathway to Canada Target 1 through the creation of Indigenous Protected and Conserved Areas in the spirit and practice of reconciliation.

Jaeger, J. A. G. (2000). Landscape division, splitting index, and effective mesh size: New measures of landscape fragmentation. Landscape Ecology, 15(2), 115–130. https://doi.org/10.1023/A:1008129329289

Keeley, A. T. H., Beier, P., & Jenness, J. S. (2021). Connectivity metrics for conservation planning and monitoring. Biological Conservation, 255, 109008. https://doi.org/10.1016/j.biocon.2021.109008

Koen, E. L., Bowman, J., Sadowski, C., & Walpole, A. A. (2014). Landscape connectivity for wildlife: Development and validation of multispecies linkage maps. Methods in Ecology and Evolution, 5(7), 626–633. https://doi.org/10.1111/2041-210X.12197

Koen, E. L., Garroway, C. J., Wilson, P. J., & Bowman, J. (2010). The Effect of Map Boundary on Estimates of Landscape Resistance to Animal Movement. PLoS ONE, 5(7), e11785. https://doi.org/10.1371/journal.pone.0011785

Krosby, M., Breckheimer, I., John Pierce, D., Singleton, P. H., Hall, S. A., Halupka, K. C., Gaines, W. L., Long, R. A., McRae, B. H., Cosentino, B. L., & Schuett-Hames, J. P. (2015). Focal species and landscape “naturalness” corridor models offer complementary approaches for connectivity conservation planning. Landscape Ecology, 30(10), 2121– 2132. https://doi.org/10.1007/s10980-015-0235-z

Laurance, W. F., Carolina Useche, D., Rendeiro, J., Kalka, M., Bradshaw, C. J. A., Sloan, S. P., Laurance, S. G., Campbell, M., Abernethy, K., Alvarez, P., Arroyo-Rodriguez, V., Ashton, P., Benítez-Malvido, J., Blom, A., Bobo, K. S., Cannon, C. H., Cao, M., Carroll, R., Chapman, C., … Zamzani, F. (2012). Averting biodiversity collapse in tropical forest protected areas. Nature, 489(7415), 290–294. https://doi.org/10.1038/nature11318

Marrotte, R. R., Bowman, J., Brown, M. G. C., Cordes, C., Morris, K. Y., Prentice, M. B., & Wilson, P. J. (2017). Multi-species genetic connectivity in a terrestrial habitat network. Movement Ecology, 5(1), 1–11. https://doi.org/10.1186/s40462-017-0112-2

Maxwell, S. L., Cazalis, V., Dudley, N., Hoffmann, M., Rodrigues, A. S. L., Stolton, S., Visconti, P., Woodley, S., Kingston, N., Lewis, E., Maron, M., Strassburg, B. B. N., Wenger, A., Jonas, H. D., Venter, O., & Watson, J. E. M. (2020). Area-based conservation in the twenty-first century. Nature, 586(7828), 217–227. https://doi.org/10.1038/s41586-020-2773-z

McNeely, J. A. (1994). Protected areas for the 21st century: Working to provide benefits to society. Biodiversity and Conservation, 3(5), 390–405. https://doi.org/10.1007/BF00057797

McRae, B. H., Dickson, B. G., Keitt, T. H., & Shah, V. B. (2008). Using circuit theory to model connectivity in ecology, evolution, and conservation. Ecology, 89(10), 2712–2724. https://doi.org/10.1890/07-1861.1

Mitchell, M. G. E., Schuster, R., Jacob, A. L., Hanna, D. E. L., Dallaire, C. O., Raudsepp-Hearne, C., Bennett, E. M., Lehner, B., & Chan, K. M. A. (2021). Identifying key ecosystem service providing areas to inform national-scale conservation planning. Environmental Research Letters, 16(1). https://doi.org/10.1088/1748-9326/abc121

Moola, F., & Roth, R. (2019). Moving beyond colonial conservation models: Indigenous Protected and Conserved Areas offer hope for biodiversity and advancing reconciliation in the Canadian boreal forest 1. Environmental Reviews, 27(2), 200–201. https://doi.org/10.1139/er-2018-0091

Naidoo, R., Balmford, A., Costanza, R., Fisher, B., Green, R. E., Lehner, B., Malcolm, T. R., & Ricketts, T. H. (2008). Global mapping of ecosystem services and conservation priorities. Proceedings of the National Academy of Sciences, 105(28), 9495–9500. https://doi.org/10.1073/pnas.0707823105

Naidoo, R., Gerkey, D., Hole, D., Pfaff, A., Ellis, A. M., Golden, C. D., Herrera, D., Johnson, K., Mulligan, M., Ricketts, T. H., & Fisher, B. (2019). Evaluating the impacts of protected areas on human well-being across the developing world. Science Advances, 5(4), eaav3006. https://doi.org/10.1126/sciadv.aav3006

Naughton-Treves, L., Holland, M. B., & Brandon, K. (2005). The role of protected areas in conserving biodiversity and sustaining local livelihoods. Annual Review of Environment and Resources, 30(1), 219–252. https://doi.org/10.1146/annurev.energy.30.050504.164507

Newmark, W. D., Halley, J. M., Beier, P., Cushman, S. A., McNeally, P. B., & Soulé, M. E. (2023). Enhanced regional connectivity between western North American national parks will increase persistence of mammal species diversity. Scientific Reports, 13(1), 474. https://doi.org/10.1038/s41598-022-26428-z

Noon, M. L., Goldstein, A., Ledezma, J. C., Roehrdanz, P. R., Cook-Patton, S. C., Spawn-Lee, S. A., Wright, T. M., Gonzalez-Roglich, M., Hole, D. G., Rockström, J., & Turner, W. R. (2021). Mapping the irrecoverable carbon in Earth’s ecosystems. Nature Sustainability, 5(1), 37–46. https://doi.org/10.1038/s41893-021-00803-6

Noss, R. F., Dobson, A. P., Baldwin, R., Beier, P., Davis, C. R., Dellasala, D. A., Francis, J., Locke, H., Nowak, K., Lopez, R., Reining, C., Trombulak, S. C., & Tabor, G. (2012). Bolder Thinking for Conservation. Conservation Biology, 26(1), 1–4. https://doi.org/10.1111/j.1523-1739.2011.01738.x

Obbard, M. E., Newton, E. J., Potter, D., Orton, A., Patterson, B. R., & Steinberg, B. D. (2017). Big enough for bears? American black bears at heightened risk of mortality during seasonal forays outside Algonquin Provincial Park, Ontario. Ursus, 28(2), 182–194. https://doi.org/10.2192/URSU-D-16-00021.1

O’Brien, P., Gunn, J. S., Clark, A., Gleeson, J., Pither, R., & Bowman, J. (2023). Integrating carbon stocks and landscape connectivity for nature-based climate solutions. Ecology and Evolution, 13(1). https://doi.org/10.1002/ece3.9725

Parks, S. A., Holsinger, L. M., Abatzoglou, J. T., Littlefield, C. E., & Zeller, K. A. (2023). Protected areas not likely to serve as steppingstones for species undergoing climate-induced range shifts. Global Change Biology, gcb.16629. https://doi.org/10.1111/gcb.16629

Parmesan, C. (2006). Ecological and Evolutionary Responses to Recent Climate Change. Annual Review of Ecology, Evolution, and Systematics, 37(1), 637–669. https://doi.org/10.1146/annurev.ecolsys.37.091305.110100

Phillips, P., Clark, M. M., Baral, S., Koen, E. L., & Bowman, J. (2021). Comparison of methods for estimating omnidirectional landscape connectivity. Landscape Ecology, 36(6), 1647– 1661. https://doi.org/10.1007/s10980-021-01254-2

Pimm, S. L., Dollar, L., & Bass, O. L. (2006). The genetic rescue of the Florida panther. Animal Conservation, 9(2), 115–122. https://doi.org/10.1111/j.1469-1795.2005.00010.x

Pither, R., O’Brien, P., Brennan, A., Hirsh-Pearson, K., & Bowman, J. (2023). Predicting areas important for ecological connectivity throughout Canada. PLOS ONE, 18(2), e0281980. https://doi.org/10.1371/journal.pone.0281980

Poley, L. G., Schuster, R., Smith, W., & Ray, J. C. (2022). Identifying differences in roadless areas in Canada based on global, national, and regional road datasets. Conservation Science and Practice, 4(4), e12656. https://doi.org/10.1111/csp2.12656

R Core Team. (2022). R: A language and environment for statistical computing (4.2.2). R Foundation for Statistical Computing. URL https://www.R-project.org/

Sawaya, M. A., Clevenger, A. P., & Schwartz, M. K. (2019). Demographic fragmentation of a protected wolverine population bisected by a major transportation corridor. Biological Conservation, 236, 616–625. https://doi.org/10.1016/j.biocon.2019.06.030

Schloss, C. A., Cameron, D. R., McRae, B. H., Theobald, D. M., & Jones, A. (2022). “No-regrets” pathways for navigating climate change: Planning for connectivity with land use, topography, and climate. Ecological Applications, 32(1), e02468. https://doi.org/10.1002/eap.2468

Schuster, R., Germain, R. R., Bennett, J. R., Reo, N. J., & Arcese, P. (2019). Vertebrate biodiversity on indigenous-managed lands in Australia, Brazil, and Canada equals that in protected areas. Environmental Science and Policy, 101, 1–6. https://doi.org/10.1016/j.envsci.2019.07.002

Sothe, C., Gonsamo, A., Arabian, J., Kurz, W. A., Finkelstein, S. A., & Snider, J. (2022). Large Soil Carbon Storage in Terrestrial Ecosystems of Canada. Global Biogeochemical Cycles, 36(2), e2021GB007213. https://doi.org/10.1029/2021GB007213

Spanowicz, A. G., & Jaeger, J. A. G. (2019). Measuring landscape connectivity: On the importance of within-patch connectivity. Landscape Ecology, 34(10), 2261–2278. https://doi.org/10.1007/s10980-019-00881-0

Spencer, W. D., Beier, P., Penrod, K., Winters, K., Paulman, C., Rustigian-Romsos, H., Strittholt, J., Parisi, M., & Pettler, A. (2010). California Essential Habitat Connectivity Project: A Strategy for Conserving a Connected California. Prepared for California Department of Transportation, California Department of Fish and Game, and Federal Highways Administration. http://www.scwildlands.org/reports/CaliforniaEssentialHabitatConnectivityProject.pdf

Taylor, P. D., Fahrig, L., Henein, K., & Merriam, G. (1993). Connectivity Is a Vital Element of Landscape Structure. Oikos, 68(3), 571. https://doi.org/10.2307/3544927

Theobald, D. M., Keeley, A. T. H., Laur, A., & Tabor, G. (2022). A simple and practical measure of the connectivity of protected area networks: The PRONET metric. Conservation Science and Practice, 4(11). https://doi.org/10.1111/csp2.12823

Theobald, D. M., Reed, S. E., Fields, K., & Soulé, M. (2012). Connecting natural landscapes using a landscape permeability model to prioritize conservation activities in the United States: Connecting natural landscapes. Conservation Letters, 5(2), 123–133. https://doi.org/10.1111/j.1755-263X.2011.00218.x

Thompson, P. L., Rayfield, B., & Gonzalez, A. (2017). Loss of habitat and connectivity erodes species diversity, ecosystem functioning, and stability in metacommunity networks. Ecography, 40(1), 98–108. https://doi.org/10.1111/ecog.02558

Tischendorf, L., & Fahrig, L. (2000). On the usage and measurement of landscape connectivity. Oikos, 90(1), 7–19. https://doi.org/10.1034/j.1600-0706.2000.900102.x

Tucker, M. A., Böhning-Gaese, K., Fagan, W. F., Fryxell, J. M., Van Moorter, B., Alberts, S. C., Ali, A. H., Allen, A. M., Attias, N., Avgar, T., Bartlam-Brooks, H., Bayarbaatar, B., Belant, J. L., Bertassoni, A., Beyer, D., Bidner, L., van Beest, F. M., Blake, S., Blaum, N., … Mueller, T. (2018). Moving in the Anthropocene: Global reductions in terrestrial mammalian movements. Science, 359(6374), 466–469. https://doi.org/10.1126/science.aam9712

Ward, M., Saura, S., Williams, B., Ramírez-Delgado, J. P., Arafeh-Dalmau, N., Allan, J. R., Venter, O., Dubois, G., & Watson, J. E. M. (2020). Just ten percent of the global terrestrial protected area network is structurally connected via intact land. Nature Communications, 11(1). https://doi.org/10.1038/s41467-020-18457-x

Watson, J. E. M., Dudley, N., Segan, D. B., & Hockings, M. (2014). The performance and potential of protected areas. Nature, 515(7525), 67–73. https://doi.org/10.1038/nature13947

Williams, D. R., Rondinini, C., & Tilman, D. (2022). Global protected areas seem insufficient to safeguard half of the world’s mammals from human-induced extinction. Proceedings of the National Academy of Sciences, 119(24), e2200118119. https://doi.org/10.1073/pnas.2200118119

