## Supplemental Table 1 for "Using sentinel nodes to evaluate changing connectivity in a protected areas network"

**Supplementary Material**

**Table S1.** Node isolation values for the 50 sentinel nodes used to evaluate connectivity of the Ontario protected areas network. Node isolation was calculated for each sentinel node as the mean pairwise effective resistance between a focal node and all other nodes. Values represent the degree of isolation for a given node, such that higher values indicate a higher degree of isolation.

| Node Park Name | Node Isolation Value (ohms) | Easting | Northing |
|---|---|---|---|
| Caliper Lake Provincial Park | 54.3861 | 6051056 | 1438843 |
| Fushimi Lake Provincial Park | 46.11147 | 6768694 | 1555866 |
| Mattagami River Beach and Aeolian Deposit Provincial Park | 46.36899 | 6885983 | 1602648 |
| Potholes Provincial Park | 46.86852 | 6769221 | 1346422 |
| Wakami Lake Provincial Park | 45.59428 | 6882357 | 1308968 |
| Little White River Provincial Park | 46.0873 | 6881164 | 1206625 |
| Driftwood Provincial Park | 83.83662 | 7286174 | 1234529 |
| Voyageur Provincial Park | 134.3989 | 7564269 | 1232139 |
| Dupont Provincial Park | 178.374 | 7526765 | 1147239 |
| Silver Lake Provincial Park | 55.08701 | 7419502 | 1109299 |
| Sauble Falls Provincial Park | 95.12488 | 7053532 | 1015410 |
| Point Farms Provincial Park | 130.8602 | 7031478 | 911433.8 |
| Wheatley Provincial Park | 285.4336 | 7002649 | 709215.5 |
| John E. Pearce Provincial Park | 155.6367 | 7076643 | 780821.8 |
| Otoskwin-Attawapiskat River Provincial Park | 45.31356 | 6486050 | 1788112 |
| Ferris Provincial Park | 303.0669 | 7337432 | 1026640 |
| Mara Provincial Park | 350.1076 | 7205988 | 1033082 |

| | | | |
|---|---|---|---|
| Opasquia Provincial Park | 45.43142 | 6121098 | 1929651 |
| Sedgman Lake Provincial Park | 45.01398 | 6493304 | 1620118 |
| Forks of the Credit Provincial Park | 126.0942 | 7170659 | 937652.9 |
| Pahngwahshahshk Ohweemushkeeg | 45.13109 | 6104357 | 1775904 |
| Spanish River Provincial Park | 45.61533 | 6974041 | 1237518 |
| Gravel River Provincial Park | 45.94743 | 6501218 | 1430700 |
| Darlington Provincial Park | 279.24 | 7268132 | 962738.8 |
| MacLeod Provincial Park | 112.548 | 6557276 | 1519359 |
| Sandbar Lake Provincial Park | 48.05959 | 6221945 | 1484763 |
| South Bay Provincial Park | 49.29436 | 7152699 | 1199690 |
| Kaiashk Provincial Park | 46.11516 | 6376026 | 1492981 |
| Chapleau-Nemegosenda River Provincial Park | 45.36905 | 6862210 | 1406958 |
| Matawin River Provincial Park | 49.00877 | 6325874 | 1362591 |
| Greenwater Provincial Park | 46.64419 | 6966264 | 1516863 |
| Lake Superior Provincial Park | 48.20546 | 6742306 | 1303496 |
| Pipestone River Provincial Park | 45.15213 | 6313211 | 1812277 |
| Weeskayjahk Ohtahzhoganeeng | 45.42276 | 6020129 | 1729357 |
| Pakwash Provincial Park | 46.73445 | 6079202 | 1622978 |
| Wabakimi Provincial Park | 44.87325 | 6371134 | 1626920 |
| Esker Lakes Provincial Park | 47.17846 | 7086520 | 1434806 |
| Winisk River Provincial Park | 46.22844 | 6508593 | 1991758 |
| Severn River Provincial Park | 45.74091 | 6272040 | 1990276 |

| | | | |
|---|---|---|---|
| French River Provincial Park | 49.7954 | 7051845 | 1165393 |
| Grassy River-Mond Lake Lowlands and Ferris Lake Uplands Provincial Park | 45.51687 | 6999711 | 1365394 |
| Sturgeon River Provincial Park | 45.87749 | 7069775 | 1280626 |
| Polar Bear Provincial Park | 45.72814 | 6758396 | 2107719 |
| Batchawana River Provincial Park | 46.19281 | 6789319 | 1267578 |
| Little Current River Provincial Park | 45.42101 | 6560062 | 1617161 |
| Obatanga Provincial Park | 47.36068 | 6702648 | 1379825 |
| St. Raphael Provincial Park | 45.21865 | 6284625 | 1619216 |
| Algonquin Provincial Park | 46.99743 | 7242207 | 1174551 |
| Short Hills Provincial Park | 188.2714 | 7244693 | 867809.1 |
| Kesagami Provincial Park | 45.52519 | 7026763 | 1688081 |
